# Context Free and Context-Dependent Conceptual Representation in the Brain

**DOI:** 10.1101/2021.05.03.442424

**Authors:** Zhiyao Gao, Li Zheng, André Gouws, Katya Krieger-Redwood, Xiuyi Wang, Dominika Varga, Jonathan Smallwood, Elizabeth Jefferies

## Abstract

How concepts are coded in the brain is a core issue in cognitive neuroscience. Studies have focused on how individual concepts are processed, but the way in which conceptual representation changes to suit the context is unclear. We parametrically manipulated the association strength between words, presented in pairs one word at a time using a slow event-related fMRI design. We combined representational similarity analysis and computational linguistics to probe the neurocomputational content of these trials. Individual word meaning was maintained in supramarginal gyrus (associated with verbal short-term memory) when items were judged to be unrelated, but not when a linking context was retrieved. Context-dependent meaning was instead represented in left lateral prefrontal gyrus (associated with controlled retrieval), angular gyrus and ventral temporal lobe (regions associated with integrative aspects of memory). Analyses of informational connectivity, examining the similarity of activation patterns across trials between sites, showed that control network regions had more similar multivariate responses across trials when association strength was weak, reflecting a common controlled retrieval state when the task required more unusual associations. These findings indicate that semantic control and representational sites amplify contextually-relevant meanings in trials judged to be related.

## Introduction

The question of how concepts are coded in the brain is a core issue in cognitive neuroscience. Neuropsychological, neuroimaging and neuromodulation studies have provided information about how individual concepts are represented in the brain (Martin 2007; Patterson et al. 2007; Binder and Desai 2011; Pulvermüller 2013; Yee et al. 2014; Lambon Ralph et al. 2017; Jefferies et al. 2020) – yet the brain produces diverse patterns of semantic retrieval for the same inputs to suit the context. For example, APPLE is associated with CAKE when it occurs together with KITCHEN, but also with LAPTOP when we encounter it with KEYBOARD. Even though concepts are thought to be constructed in this dynamic fashion, empirical studies have, until recently, largely focused on invariant conceptual representation – i.e. the features of concepts that do not vary across contexts (Yee and Thompson-Schill 2016). We therefore presented thematically related word-pairs which varied from weak to strong associations to instantiate context-dependent representations of concepts, to investigate the neural basis of flexible semantic cognition (Yee and Thompson-Schill 2016).

The controlled semantic cognition (CSC) framework suggests that distributed modality-specific features (e.g. visual, auditory, motor and valence features) in ‘spoke’ systems are integrated within a semantic ‘hub’ or ‘convergence zone’ in the anterior temporal lobes (ATL), giving rise to heteromodal concepts (Patterson et al. 2007; Lambon Ralph et al. 2017). An additional distributed semantic control network manipulates activation within this conceptual representation system to generate appropriate patterns of semantic retrieval that suit the circumstances in which they occur. In well-practiced contexts, left angular gyrus (AG) and ATL are thought to support conceptual combination, with the strongest responses observed when conceptual retrieval is highly coherent and control demands are minimized (Bemis and Pylkkänen 2013; Davey et al. 2015; Teige et al. 2019; Lanzoni et al. 2020). In other situations, when retrieval must be focused on non-dominant features or unusual conceptual combinations, there is greater engagement of the ‘semantic control network’, which includes left inferior frontal gyrus (IFG) (Thompson-Schill et al. 1997; Wagner et al. 2001; Whitney et al. 2011; Hallam et al. 2016; Hallam et al. 2018; Gonzalez Alam et al. 2019; Lanzoni et al. 2020; Jackson 2021). These semantic control processes can shape the interaction between hub and spokes to focus on the features required by a task (Davey et al. 2016; Lambon Ralph et al. 2017; Chiou et al. 2018; Zhang et al. 2021). Stronger connectivity between left IFG and the semantic ‘hub’ region in left ventral ATL is associated with better semantic controlled semantic cognition (Chiou and Lambon Ralph 2019; Jung et al. 2021). Therefore, we reasoned that the semantic control network is a neural candidate underlying context-dependent meaning.

In some situations, there is an explicit goal for semantic retrieval specified by the task demands: for example, for the concept ‘PIANO’, if we want to play this instrument, our retrieval is focused on the motor features that allow us to move our fingers in an appropriate way, while if we have the goal of finding this instrument in a warehouse, we will retrieve visual information about its shape and size. In these situations, semantic control processes might be able to bias the pattern of semantic retrieval in task-appropriate ways by facilitating or inhibiting connections between the heteromodal hub in ATL and task-relevant and task-irrelevant spokes. Multivoxel pattern analysis (MVPA) provides us with a powerful tool to probe how the representation of semantic information in the brain varies according to the context; these studies have started to explore how features combine to construct concepts and how word meaning is modified syntactically (Allen et al. 2012; Coutanche and Thompson-Schill 2014; Boylan et al. 2015; Hoffman and Tamm 2020; Solomon and Thompson-Schill 2020). For example, a recent magnetoencephalography study showed that neural representations of the noun were modified across temporal, inferior frontal and inferior parietal regions according to the verb it was combined with (Lyu et al. 2019). Yet in many other situations requiring semantic control – for example, when weak as opposed to strong thematic associations must be identified – participants are not required to focus on specific types of features, but instead to identify a context in which concepts co-occur. Given that there is no explicit goal or instruction guiding semantic retrieval, this might require participants to create an event representation to simulate or construct a scenario, which can then bias retrieval towards features of the concept that are consistent with this event, and away from other potentially dominant features which are inconsistent (Mirman et al. 2017). An understanding of the neurobiological mechanisms that underpin this process remains elusive.

In the current study, we used fMRI to identify where in the brain non-contextualized meanings of words are represented as well as to determine how words are integrated to form context-dependent conceptual representations. We varied the strength of thematic relationships between two words presented successively, from very strong (dog with leash), through intermediate trials (dog with beach) to very weak pairs (dog with keyboard). We leveraged word embeddings of natural language processing (NLP) to establish vectors of similarity for our word stimuli which were either (i) focused on context-invariant meaning using word2vec (Mikolov et al. 2013) or (ii) captured vectors of similarity for words based on the ongoing context (i.e. taking into account the preceding/following words) using ELMo (Peters et al. 2018). We combined these computational linguistic approaches with a slow-event related fMRI design and representational similarity analysis (RSA) (Kriegeskorte et al. 2006; Kriegeskorte et al. 2008), implemented using a searchlight approach, to determine where in the brain similarity in multivoxel activity patterns could be predicted by context-free and context-sensitive conceptual similarities. Specifically, we asked whether networks implicated in more automatic and controlled aspects of semantic cognition in previous studies (Fedorenko et al. 2013; Humphreys and Lambon Ralph 2015; Davey et al. 2016; Wang et al. 2020; Gao et al. 2021; Jackson 2021) would show differential representation of context-independent and context-dependent meaning, or alternatively whether semantic regions across these networks would commonly support the construction of context-dependent meanings but in different ways (via more automatic vs. controlled integrative processes, giving rise to context-dependent meanings of strong and weak associations respectively).

## Materials and Methods

### Participants

A group of 32 healthy participants aged 19 to 35 years (mean age = 21.97 ±3.47 years; 19 females) was recruited from the University of York. They were all right-handed, native English speakers, with normal or corrected-to-normal vision and no history of psychiatric or neurological illness. The study was approved by the Research Ethics Committee of the York Neuroimaging Centre. All volunteers provided informed written consent and received monetary compensation or course credit for their participation. Data from four participants was excluded due to head motion (translational displacement was greater than 2mm), resulting in a final sample of 28 participants for the semantic task. This study provides a novel analysis of a dataset first reported by (Gao et al. 2021).

### Semantic Task

The experimental stimuli were 192 English concrete noun word pairs. We excluded any abstract nouns and pairs of items drawn from the same taxonomic category, so that only thematic links were evaluated. The strength of the thematic link between the items varied parametrically from trials with no clear link to highly related trials; in this way, participants were free to decide based on their own experience if the words had a discernible semantic link. There were no ‘correct’ and ‘incorrect’ responses: instead, we expected slower response times and less convergence across participants for items judged to be ‘related’ when the associative strength between the items was weak, and for items judged to be ‘unrelated’ when the associative strength between the items was strong. Overall, there were roughly equal numbers of ‘related’ and ‘unrelated’ responses across participants.

Each trial began with a visually presented word (WORD-1) which lasted 1.5s, followed by a fixation cross presented at the centre of the screen for 1.5s. Then, the second word (WORD-2) was presented for 1.5s, followed by a blank screen for 1.5s. Participants had 3s from the onset of WORD-2 to judge whether this word pair was semantically associated or not by pressing one of two buttons with their right hand (using their index and middle fingers). During the inter-trial interval (3s), a red fixation cross was presented until the next trial began. Both response time (RT) and response choice were recorded. Participants finished 4 runs of the semantic task, each lasting 7.3 min. Before the scan, they completed a practice session to familiarise themselves with the task and key responses.

### Neuroimaging Data Acquisition

Imaging data were acquired on a 3.0 T GE HDx Excite Magnetic Resonance Imaging (MRI) scanner using an eight-channel phased array head coil at the York Neuroimaging Centre. A single-shot T2*-weighted gradient-echo, EPI sequence was used for functional imaging acquisition with the following parameters: TR/TE/θ = 1500 ms/15 ms/90°, FOV = 192 × 192 mm, matrix = 64 × 64, and slice thickness = 4 mm. Thirty-two contiguous axial slices, tilted upper to the eye, were obtained to decrease distortion in the anterior temporal lobe and prefrontal cortex. Anatomical MRI was acquired using a T1-weighted, 3D, gradient-echo pulse-sequence (MPRAGE). The parameters for this sequence were as follows: TR/TE/θ = 7.8s/2.3 ms/20°, FOV = 256 × 256 mm, matrix = 256 × 256, and slice thickness = 1 mm. A total of 176 sagittal slices were acquired to provide high-resolution structural images of the whole brain.

### Semantic Similarity Matrices

Using natural language processing tools, two semantic similarity matrices were constructed based on two types of word embedding to investigate different types of semantic information in neural activity patterns. Embedding vectors extracted from word2vec and ELMo for all word pairs are available online: https://osf.io/hwfdp/.

### word2vec

The word2vec model represents words as fixed high-dimensional vectors of embeddings. The vectors of word embeddings were generated by training the network on the 100-billion-word Google News corpus. Each time the network was presented with a word from the corpus, it was trained to predict the context in which it appeared, where context was defined as the two words preceding and following it in the corpus. The model learns to represent words used in similar contexts with similar patterns; each word’s vector had 300 dimensions, with similarity across two words’ vectors indicating that they appear in similar contexts, and thus have related meanings. Word2vec embeddings are fixed and unique for each word; for example, irrespective of whether ‘apple’ was followed by ‘bread’ or ‘keyboard’, its word embeddings were the same. Therefore, using word2vec, we constructed semantic similarity matrices (word2vec-based RSM), separately for WORD1 and WORD2; these reflected the meaning of single words, unmodified by the context in which these items appeared, by calculating cosine similarity between words drawn from different trials.

### ELMo

Given that context can change the meaning of individual words in sentences and phrases, Peters et al. (2018a) proposed a deep contextualized word embedding model called ELMo (Embeddings from Language Models) to capture the context-dependent semantic representation of words. Rather than providing a dictionary of words and their corresponding vectors, ELMo analyses words within their linguistic context, with each token assigned a representation that is a function of the entire input sentence. ELMo representations are deep in the sense that they are a function of all the internal layers of a deep bidirectional language model: there is a context-independent fixed input vector for the word in the lowest layer, with two higher layers capturing backward and forward context-sensitive aspects of word meaning. We used the pretrained model released by Allennlp (Gardner et al. 2018), which was trained on a large test corpus of 5.5B tokens from Wikipedia and the English news data from the workshop of machine translation (WMT) 2008-2012. We selected the top layer in ELMo to generate context-sensitive embeddings for WORD2. Each vector representing word meaning had 1024 dimensions. We calculated a context-sensitive semantic similarity matrix (ELMo-based RSM) for WORD-2 by correlating the top embedding vectors across words taken from different trials, regressing out the lowest layer’s embedding vectors to control the contribution of more context-independent patterns of representation (see Xu et al. (2018) for a similar approach), to search for brain regions where the pattern of responses across voxels was associated with contextually-constrained semantic cognition.

To further validate this approach, we searched for sentences that included the word pairs used in the current study (within widely used NLP datasets, such as Google News) and estimated the context-dependent meaning of WORD-2 stimuli within these sentence contexts (see Supplementary Materials). Following the procedure described above, we constructed a sentence-based context-dependent meaning similarity matrix. The two similarity matrices for context-dependent meaning were strongly correlated; r = 0.81 (p < 0.001) and the neuroimaging results are also highly consistent (see Supplementary Materials).

### fMRI Data Preprocessing Analysis

Image preprocessing and statistical analysis were performed using FEAT (FMRI Expert Analysis Tool) version 6.00, part of FSL (FMRIB software library, version 5.0.9, www.fmrib.ox.ac.uk/fsl). The first 4 volumes before the task were discarded to allow for T1 equilibrium. The remaining images were then realigned to correct for head movement. Translational movement parameters never exceeded 1 voxel in any direction for any participant or session. No spatial smoothing was performed. The data were filtered in the temporal domain using a nonlinear high-pass filter with a 100s cutoff. Following Deuker et al. (2016); Bellmund et al. (2019), six motion parameters were used as predictors in a GLM. The residuals from this model (which could not be explained by motion) were then taken into the next analysis step. A two-step registration procedure was used whereby EPI images were first registered to the MPRAGE structural image (Jenkinson and Smith 2001). Registration from MPRAGE structural image to standard space was further refined using FNIRT nonlinear registration (Andersson et al. 2007, 2007). The denoised time series were transformed to standard space for the multivariate analyses.

### Univariate Parametric Analysis

We examined the effects of semantic control demands via a parametric manipulation of strength of association at the network level, following the approach reported by Gao et al. (2021). We predicted that it would be harder for participants to decide that items were semantically related when they were weakly associated (with lower word2vec values), and it would also be harder for them to decide that items were semantically unrelated when in trials with higher word2vec values. Therefore, we extracted the parametric effect of word2vec on the BOLD response separately for trials judged to be related and unrelated. Since association strength was negatively correlated with control demands for trials judged to be related, we means-centered and reversed the sign of word2vec values for these trials in each run before the next analysis step. This allowed us to compare the effects of semantic control demands across related and unrelated trials. We performed this analysis within four functional networks involved in more automatic or more controlled aspects of semantic cognition or executive control. The networks were taken from previous meta-analytic studies of the semantic control network (SCN) and multiple-demand network (MDN) (Fedorenko et al. 2013; Jackson 2021). Within these networks, we selected 1) semantic control specific areas, which did not overlap with MDN; 2) multiple-demand specific regions, which did not overlap with SCN; 3) shared control regions, identified from the overlap between MDN and SCN; and 4) semantic regions not implicated in control; these were identified using Neurosynth (search term ‘semantic’; 1031 contributing studies; http://www.neurosynth.org/analyses/terms/), removing regions that overlapped with the two control networks to identify regions associated with semantic representation or more automatic aspects of semantic retrieval, mostly within DMN (e.g. in lateral temporal cortex and angular gyrus). This process defined thirty ROIs; four in semantic non-control areas, three in SCN, six in the overlap of MDN and SCN, and seventeen in MDN specific areas. These thirty ROIs are available online: https://osf.io/hwfdp/ and were previously used by Gao et al. (2021). The ROIs within each network were averaged across all relevant sites for the network-based analyses presented below.

### Pattern Similarity Analysis

In order to examine how the characteristics of semantic representation were influenced by the context, we focused on the decision phase of the task. This period corresponded to TR 6 and 7 after WORD-1 onset. Second-order representational similarity analysis (RSA) was performed using a searchlight approach; semantic RSMs (i.e., the word2vec-based RSM and ELMo-based RSM) were compared with neural pattern similarity matrices (brain-based RSM) to test what semantic information was represented in different brain regions. Neural pattern similarity was estimated for cubic regions of interest (ROIs) containing 125 voxels surrounding a central voxel, as many previous studies examining semantic representation used this approach successfully (Fairhall and Caramazza 2013; Malone et al. 2016; Stolier and Freeman 2016; Leshinskaya et al. 2017; Wang et al. 2017; Viganò and Piazza 2020). In each of these ROIs, we compared patterns of brain activity to derive a neural RSM from the pairwise Pearson correlations of each pair of trials. We excluded any pairs presented in the same run from the calculation of pattern similarity to avoid any auto-correlation issues. Spearman’s rank correlation was used to measure the alignment between semantic and brain-based models during the decision phase. Of note, both semantic models (word2vec and ELMo-based RSMs) were correlated to the same neural similarity matrices, which allows us to examine where and how context-dependent and context-free meanings of concepts were represented in the brain, depending on the decision participants reached (i.e., related versus unrelated) during the decision phase. The resulting coefficients were Fisher’s z transformed and statistically inferred across participants. The searchlight analysis was conducted in standard space. A randomeffects model was used for group analysis. Since no first-level variance was available, an ordinary least square (OLS) model was used.

We also examined neural representations of context-free and context-dependent meaning within regions of interest (ROIs). As for the univariate analysis of parametric effects of word2vec, we focused on four sets of regions: 1) semantic control specific (SCN specific) areas, which did not overlap with MDN; 2) multiple-demand specific (MDN specific) regions, which did not overlap with SCN; 3) shared control regions identified from the overlap between MDN and SCN; and 4) regions within the semantic network not implicated in control. The same thirty ROIs were used for both univariate and multivariate analyses, with individual ROIs within each network averaged for network-based analyses.

### Informational Connectivity between Networks as a Function of Association Strength

Even when multiple networks show similar representation of context-dependent meanings based on second-order RSA (ELMo to neural alignment), this does not establish that they represent similar information across trials. In order to examine whether neural activity patterns between regions belonging to specific functional networks capture similar semantic representations, and to investigate how this similarity in the multivariate response across trials might change as a function of the strength of association between the words being linked, we performed a novel informational connectivity analysis. In contrast to functional connectivity analysis using global BOLD signals averaged across voxels in each region, this analysis assessed the similarity of the multivariate patterns between pairs of brain regions across trials (Aly and Turk-Browne 2016; Xiao et al. 2017; Anzellotti and Coutanche 2018), within sliding windows capturing trials of different associative strengths. First, we sorted all the word-pairs from weakly to strongly associated according to their semantic association strength (word2vec value) for the related and unrelated conditions separately. Next, we grouped every 16 trials into one window; adjacent windows partially overlapped with each other by 4 trials. We then computed second-order RSAs by correlating the neural similarity matrices between ROIs within each window. The next step of this analysis established how this informational connectivity metric changed as a function of the association strength of the words being linked, using Spearman correlation. The resulting correlation coefficients were transformed into Fisher’s z-scores and then averaged across ROIs within each network. We performed several variants of this analysis, using window sizes and overlapping step sizes of 16,4; 12,4; 20,4, respectively, (window sizes, i.e., the number of trials in each window varying in associative strength; step sizes, i.e., the number of overlapping trials across adjacent windows), to ensure the robustness of our conclusions.

### Mixed-Effects Modelling Analysis of Behavioural Performance

Since participants judged different numbers of items to be semantically related and unrelated, mixed-effects modelling was used for the analysis of the behavioural data. This approach is particularly suitable when the number of trials in each condition differs across participants (Mumford and Poldrack 2007; Ward et al. 2013). Mixed-effects modelling was implemented with lme4 in R (Bates et al. 2014). We used the likelihood ratio test (i.e., Chi-Square test) to compare models, in order to determine whether the inclusion of predictor variables significantly improved the model fit. Semantic association strength was used as a predictor of the decision participants made (judgements about whether the words were related or unrelated) and, in a separate model, the reaction time this decision took. Participant identity was included as a random effect. By comparing models with and without the association strength predictor, we were able to establish whether semantic association strength predicted semantic performance.

## Results

### Behavioural Results

Since we used a continuous manipulation of associative strength, and there is no categorical boundary of word2vec values which can capture the trials reliably judged to be related and unrelated, traditional error scores were not calculated. Chi-square was conducted to examine whether equal numbers of word pairs were judged to be related or unrelated by the participants (mean ratio for the related and unrelated trials: 0.491 vs. 0.495, χ2(1) = 0.00021, p > 0.995). Linear mixed-effects model analysis revealed that the strength of the semantic association (word2vec value for each pair) was positively associated with a higher probability that participants would identify a semantic relationship between the words (χ2(1) = 2505.4, p < 0.001). The percentage of trials judged to be related varied from 34.9% to 60.9% with a standard deviation of 6.16%, while the percentage of trials judged to unrelated ranged from 39.1 to 63.5% with a standard deviation of 6.16%. There were no outliers in these judgements of relatedness (no participants were more than 3 standard deviations from the mean).

Linear mixed-effects models also examined how association strength modulated reaction time (RT) for trials judged to be related and unrelated. There was a significant effect of strength of semantic association (word2vec) for both related and unrelated decisions: association strength was negatively associated with RT for related trials (χ2(1) = 156.55, p = 2.2e-16), and positively associated with RT for trials judged to be unrelated (χ2(1) = 52.415, p =4.5e-13); Figure 1B. It was more difficult for participants to retrieve a semantic connection between two words when the strength of association was lower; on the contrary, it was easier for them to decide there was no semantic connection between word pairs with when word2vec was low. The average reaction time for trials judged to be related was 1.12s (standard deviation = 0.48s), while the average time for unrelated judgements was 1.17s (standard deviation = 0.47s). 0.9% and 0.7% of related and unrelated decisions respectively were outliers (more than 3 standard deviations from the mean).

**Figure 1.**
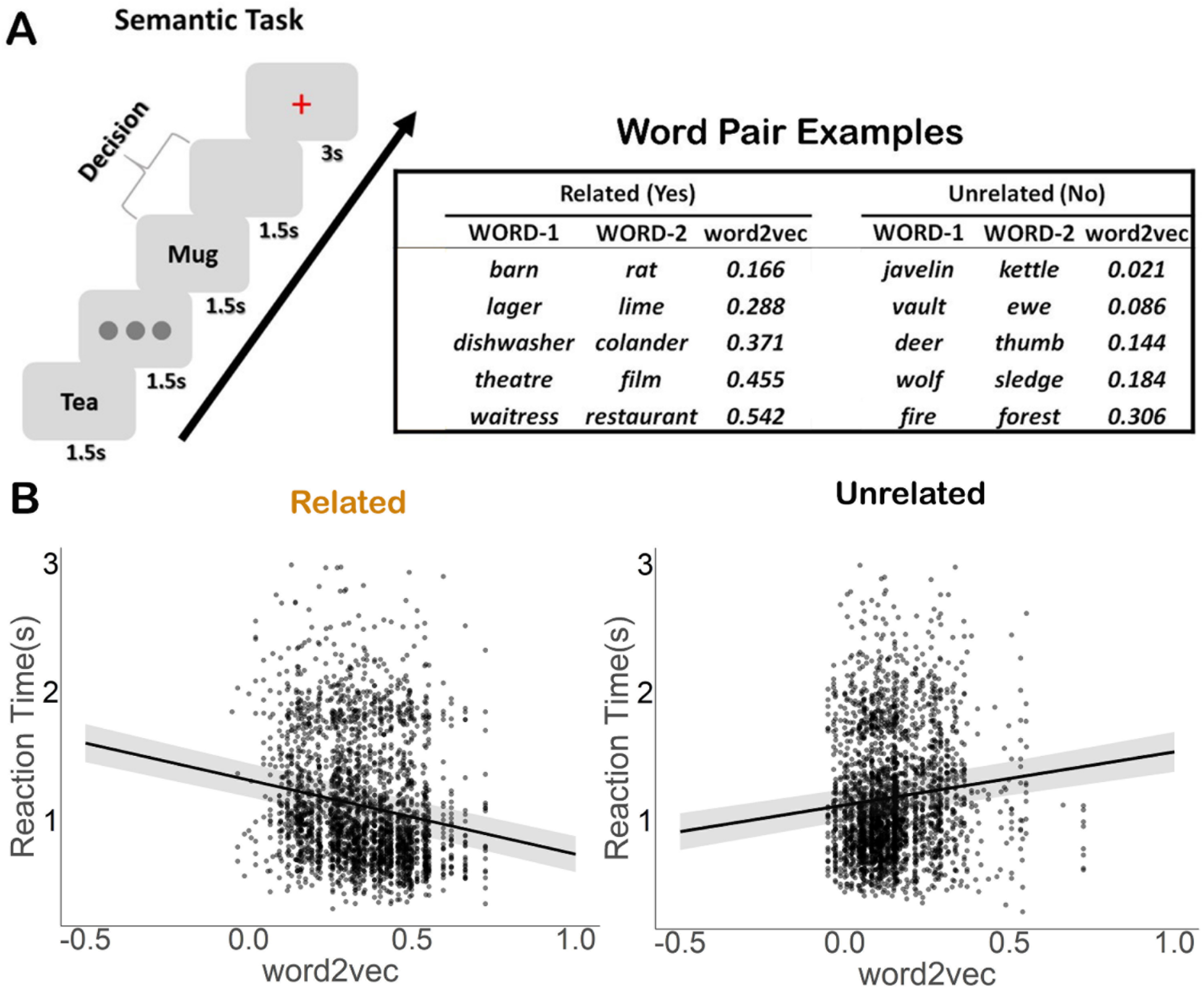
Experiment paradigm and behavioural results. A. Left-hand panel: Semantic association task; participants were asked to decide if word pairs were semantically related or not. Right-hand panel: Word pair examples for both related and unrelated decisions from one participant, with association strength increasing from weak to strong. Trials were assigned to related and unrelated sets on an individual basis for each participant, depending on their decisions. B. The semantic association strength (word2vec) was negatively associated with reaction times for related trials and positively associated with reaction time for trials judged to be unrelated. People were faster to discern a relationship between words when they had high semantic overlap, and slower to decide that the words were unrelated when they had high semantic overlap.

**Figure 2.**
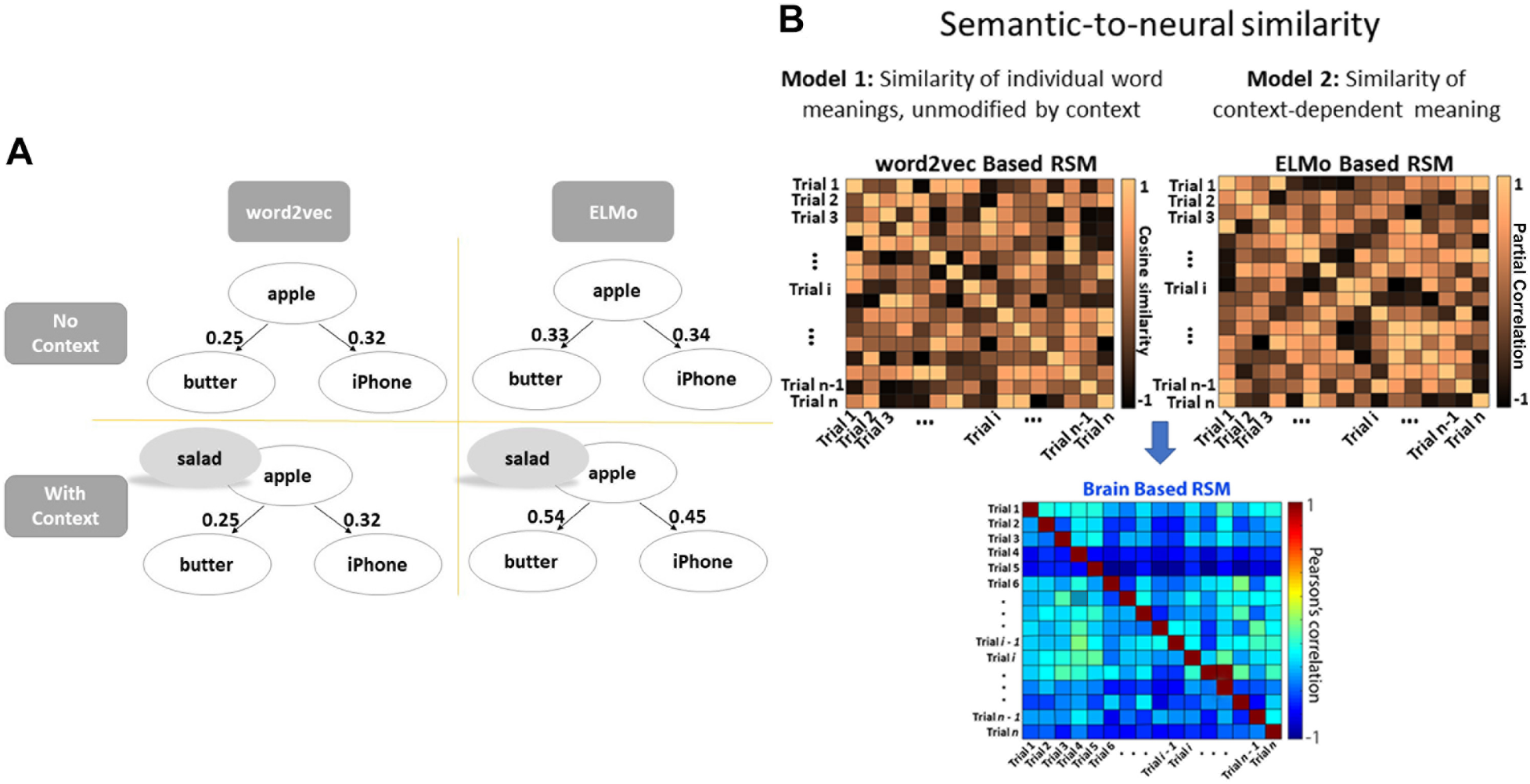
A. Example association strength values produced by ELMo and word2vec. The word2vec value between word-pairs was fixed and not dependent on the context the words were presented in, in contrast to ELMo. B. Semantic-to-neural similarity computed via second-order RSAs: these analyses characterized the semantic similarity between words on different trials and examined the association with neural pattern similarity across trials. Left-hand panel: word2vec-based RSM for unmodified word meanings across trials – this matrix captured the semantic similarity of individual words used across trials; right-hand panel: ELMo-based RSM for context-dependent meaning – this matrix captured the semantic similarity of contextually-modified meanings across trials.

### fMRI Results

#### Neural Representation of Context-Free Meaning

Whole-brain analysis was performed using a searchlight approach. First, we examined context-free semantic representation of the original or unmodified meaning of individual words during the decision phase, using the word2vec model to assess semantic similarity across trials - since word similarity in this model is fixed, and not dependent on the context in which words are presented. The strongest responses reflecting context-free meaning are expected for WORD-1, since retrieval of the meaning of this item commenced in the absence of any semantic context (while for WORD-2, the context established by the first word in the pair is likely to influence the pattern of retrieval). We also expect context-free meaning to be most relevant during trials judged to be unrelated, since on these trials, participants did not identify a linking context.

For WORD-1, on those trials judged to be semantically unrelated (i.e., when no linking context was retrieved), a significant positive association between neural pattern similarity and semantic similarity based on word2vec was seen in the left supramarginal gyrus; see Figure 3A (left-hand panel). This site showed more similar neural patterns during semantic decisionmaking when the context-free meaning of WORD-1 was more similar. For word pairs that were judged to be semantically related, there was no relationship between neural pattern similarity and semantic similarity for WORD-1.

**Figure 3.**
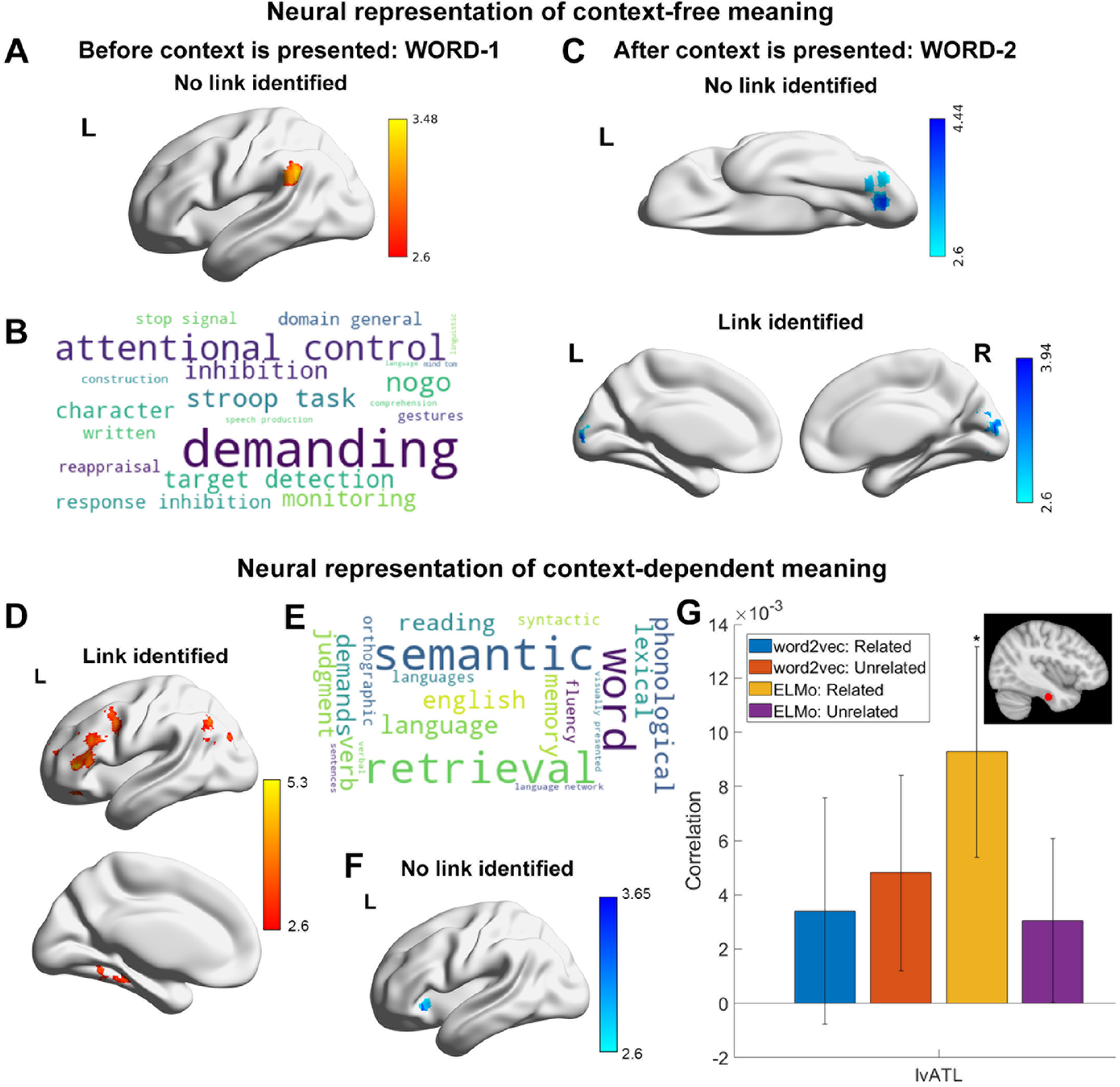
A. Positive correlation was found for the neural representation of original meaning of WORD-1 (before context is presented) on trials judged to be unrelated (Z > 2.6, corrected). B. Decoding of this cluster-corrected spatial map (A) using Neurosynth revealed terms linked to attention control and task demands. C. Negative correlations were found for the neural representation of original meaning of WORD-1 (before context is presented) and WORD-2 (after context is presented) for items judged to be related. D. Positive correlation was found for the neural representation of context-dependent meaning of WORD-2 for trials judged to be related (Z > 2.6, corrected). E. Decoding of this cluster-corrected spatial map (D) using Neurosynth revealed terms linked to semantic and language processing. F. Negative correlation was found for the neural representation of context-dependent meaning of WORD-2 for the trials judged to be unrelated (Z > 2.6, corrected). G. Region of interest analysis: A spherical ROI (117 voxels, right top panel) was created for the left ventral anterior temporal lobe (lvATL) around the peak voxel at MNI coordinate (x = −36, y = −18, z = −30) reported by Binney et al. (2016). Significant positive correlation was found for the neural representation of context-dependent meaning of WORD-2 in trials judged to be related. * p < 0.05.

Next, we examined the representation of original word meaning for WORD-2. The meaning of this item was retrieved in a semantic context established by the presentation of WORD-1, and consequently, we did not expect to see an association between neural pattern similarity and context-free meaning across trials. In line with our expectations, there were no positive correlations between the word2vec and neural models for WORD-2; instead, there were negative correlations between these models in visual cortex; see Figure 3B. These negative associations suggest that the prior presentation of WORD-1 pushed the visual representations of semantically similar WORD-2 items further apart. Semantically similar items often have similar visual features – for example, animals typically have legs and eyes; vehicles often have wheels; fruits are often brightly coloured. Our results suggest that when participants retrieve word meaning in a context established by the presentation of a previous item, they focus less on these shared visual features of semantically similar concepts.

In summary, evidence for the neural representation of context-free word meaning was only found for WORD-1 in unrelated trials. There was no evidence that participants represented context-free meanings either for WORD-2 (when participants were attempting to retrieve a semantic link with the previous word) or for trials in which the words were judged to be related in meaning, indicating that a linking context was retrieved.

### Neural Representation of Context-Dependent Meaning

The preceding results demonstrated that activity patterns in the brain represented the original or unmodified meaning of words presented in the absence of a context, but not when a linking context was retrieved. Motivated by the theory that a concept cannot be meaningfully separated from the context in which it occurs (Yee and Thompson-Schill 2016), we next tested whether neural similarity across trials was related to contextually-derived word meaning, especially for word pairs judged to be related. We focused this analysis on WORD-2, since the meaning of this item was processed in the context of the preceding item (in contrast, no semantic context was available when the meaning of WORD-1 was first retrieved). We used ELMo to estimate the context-dependent semantic similarity between the WORD-2 items across trials, separately for words presented in trials judged to be related and unrelated. For trials judged to be semantically related, a positive correlation between neural similarity and ELMo-based semantic similarity was found in left lateral frontal cortex and angular gyrus; see Figure 4A (right-hand panel). No correlations between context-dependent semantic similarity and neural similarity were found for trials judged to be unrelated. Additional analyses were conducted using a sentences-based context-dependent meaning estimation, which produced highly similar results showing a positive correlation between neural similarity and ELMo-based semantic similarity in left lateral frontal cortex and angular gyrus, see Supplementary Materials Figure S1A.

**Figure 4.**
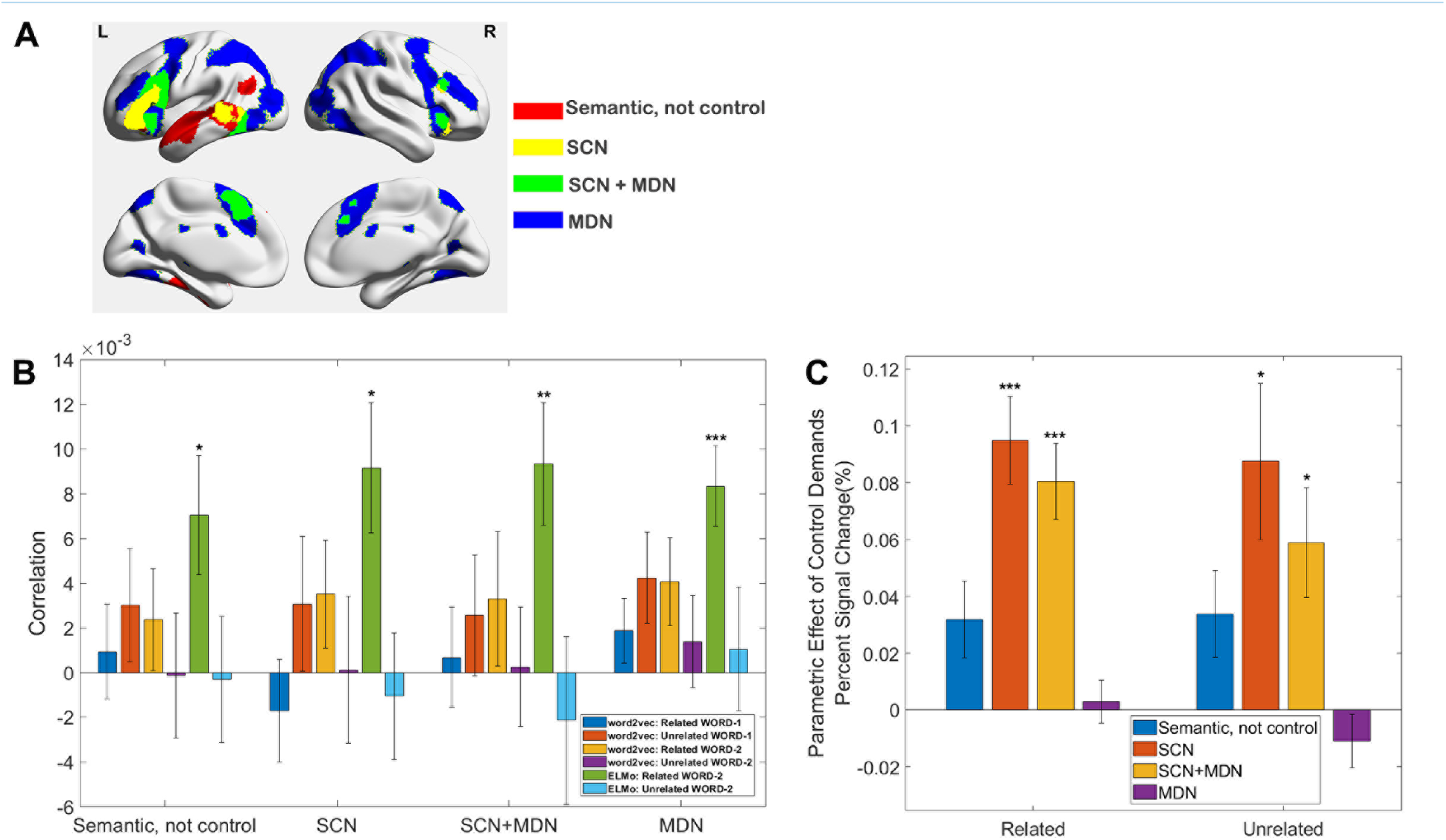
A: Functional networks: (i) semantic not control, (ii) within the semantic control network (SCN) but outside multiple-demand cortex (DMN), (iii) within both SCN and MDN, and (iv) falling in MDN regions not implicated in semantic cognition. B. Neural representation of context-free and context-dependent meaning in functional networks. Positive correlations were found for context-dependent meaning of WORD-2 for trials judged to be related in all four networks. C. Univariate parametric effects in four functional networks showing modulation of the BOLD response according to control demands: the weaker associative strength for trials judged to be related were associated with the higher activation; while the stronger associative strength for trials judged to be unrelated were associated higher activation. SCN and regions falling within both SCN and MDN showed significantly higher activation for those trials with weaker associations and consequently higher controlled retrieval demands. * p < 0.05; ** p < 0.01; *** p < 0.001. Bonferroni correction was applied.

Left ventral anterior temporal lobe (lvATL) has been suggested to be a semantic ‘hub’ (Binney et al. 2016; Lambon Ralph et al. 2017), playing a crucial role in representing strong associations and semantic combinations in long-term memory (Bemis and Pylkkänen 2013; Teige et al. 2019). However, distortion and signal loss occur in this area due to magnetic inhomogeneities close to air-tissue boundaries, causing a lower signal-to-noise ratio and weaker effects of interest (Weiskopf et al. 2006; Binney et al. 2010); see Supplementary Materials Figure S4. Given we did not observe effects in ventral ATL in whole-brain analyses, the neural representation of context-free and context-dependent meaning at this site was assessed using ROI-based analysis. We created a sphere ROI (117 voxels) for lvATL around the peak voxel implicated in semantic cognition at MNI coordinate (x = −36, y = −18, z = −30) (Binney et al. 2016). Only neural representation of context-dependent meaning for related trials was found, see Figure 3F. To check the robustness of our results, additional analyses were conducted using both larger (179 voxels) and smaller spheres (81 voxels) centered on lvATL; highly similar results were found, see Supplementary Materials Figure S1B.

### What dominates the semantic response within functional networks?

To examine how context-free and context-dependent meaning is represented in functional networks relevant to semantic representation and control, we conducted second-order RSA analyses for each ROI within four networks, reporting averages across the ROIs for each network. These functional networks included semantic not control areas (which are implicated in semantic processing but not in semantic or domain-general control), semantic control areas (i.e. cortical regions specifically implicated in semantic control and not domain-general control) and areas shared by semantic control and multiple-demand network (MDN) areas as well as areas specific to MDN that are not typically activated by semantic tasks (see more details in Methods). The results showed that there was no significant neural representation of context-free meaning in any of these networks, but there was significant representation of context-dependent meaning for WORD-2 for those trials judged to be related in all four networks. Moreover, there was no significant difference between networks in the representation of context-dependent meaning and context-free meaning for WORD-1 and WORD-2 for trials either judged to be related or unrelated (student’s T test between any two pairs, all Ps > 0.45, after Bonferroni correction), suggesting all four functional networks track the way that words are being used, instead of long-term invariant semantic knowledge.

We further examined whether the representation of context-dependent meaning was dependent on association strength across networks. We sorted trials by strength of association and grouped those trials judged to be related into small analysis ‘windows’ containing 16 trials (window length). Adjacent windows were overlapping by 4 trials (step size). We measured neural representations of context-dependent meaning in each window and correlated the neural representation with association strength using spearman correlation. The above procedure was conducted for each ROI and averaged across ROIs within each network. No significant linear relationship between association strength and the neural representation of context-dependent meaning was found in any of these functional networks (all Ps > 0.85, after Bonferroni correction).

Even though context-dependent meaning was represented irrespective of associative strength across these different networks, previous studies suggest that they are differently sensitive to semantic control demands (Fedorenko et al. 2013; Humphreys and Lambon Ralph 2015; Davey et al. 2016; Jackson 2021). To confirm this pattern in the current dataset, we characterised the parametric effects of associative strength (inverted for related trials such that higher scores denote greater activation for more difficult decisions, for both related and unrelated judgments). We conducted a two-way repeated ANOVA, with the factors of network (4 levels) and trial type (related vs. unrelated) as within-participant variables. We found a significant main effect of network (F(1.429, 38.591) = 30.737, p < 0.001) but no main effect of trial type (F(1, 27) = 0.477, p = 0.496) and no interaction (F(1.995, 53.861) = 0.877, p = 0.422); see Figure 4C. Direct comparisons between networks using t-tests revealed significantly stronger responses to the semantic difficulty manipulation in SCN than in both ‘semantic not control’ regions (p < 0.001) and MDN regions that were outside those areas activated by semantic control manipulations (p < 0.001). There was no significant difference between SCN and SCN+MDN regions (p = 0.47); SCN+MDN areas also showed significantly stronger responses to difficulty than ‘semantic not control’ regions (p < 0.001) and MDN (p < 0.001). All p values were Bonferroni corrected. These results suggest that different brain networks play distinct roles in semantic retrieval.

### Neural representation between networks was more differentiated as association strength increased for related trials

While there were no differences between networks in the neural representation of context-dependent meaning and context-free meaning in the analysis above, this does not demonstrate that these networks represent conceptual information in the same way, especially given that our univariate analysis shows different responses across these networks to the parametric manipulation of association strength. In order to assess the degree to which neural representation was similar across networks, and how this similarity in neural patterns changed as a function of association strength, we conducted a novel ‘sliding window’ analysis of informational connectivity. We firstly measured the overall informational connectivity between networks when all trials were included for related and unrelated decisions separately. No significant differences were found overall for informational connectivity between related and unrelated trials (all Ps > 0.5 after FDR correction; see Supplementary Materials Figure S2A). Next, we sorted trials judged to be related according to their associative strength, from weak to strong (based on word2vec between the words in each pair) and grouped every 16 trials into one window; we then constructed neural similarity matrices in each window by calculating the Spearman’s correlation of neural similarity matrices between pairs of ROIs, taking an average across ROIs belonging to each network. This allowed us to calculate Spearman correlation between association strength and informational connectivity at the network level. All correlation values were Fisher’s Z transformed. There was a significant effect of associative strength on informational connectivity between networks for related trials; the multivariate pattern similarity between related trials was increased when strength of association was low for the SCN+MDN regions (Figure 5B). This finding suggests that these regions take on a pattern of connectivity that supports controlled semantic retrieval; these connections are more similar across trials that are weakly related. No such effects were found for those trials judged to be unrelated. Further direct comparisons of the influence of associative strength on informational connectivity between related and unrelated trials revealed significantly faster decreases in informational connectivity for related trials as association strength increased: this pattern was observed when SCN+MDN regions were compared with SCN (p = 0.004), MDN (p = 0.008), and other SCN+MDN parcels (p = 0.005), this effect was not significant within or between any other networks. All p values were Bonferroni corrected.

**Figure 5.**
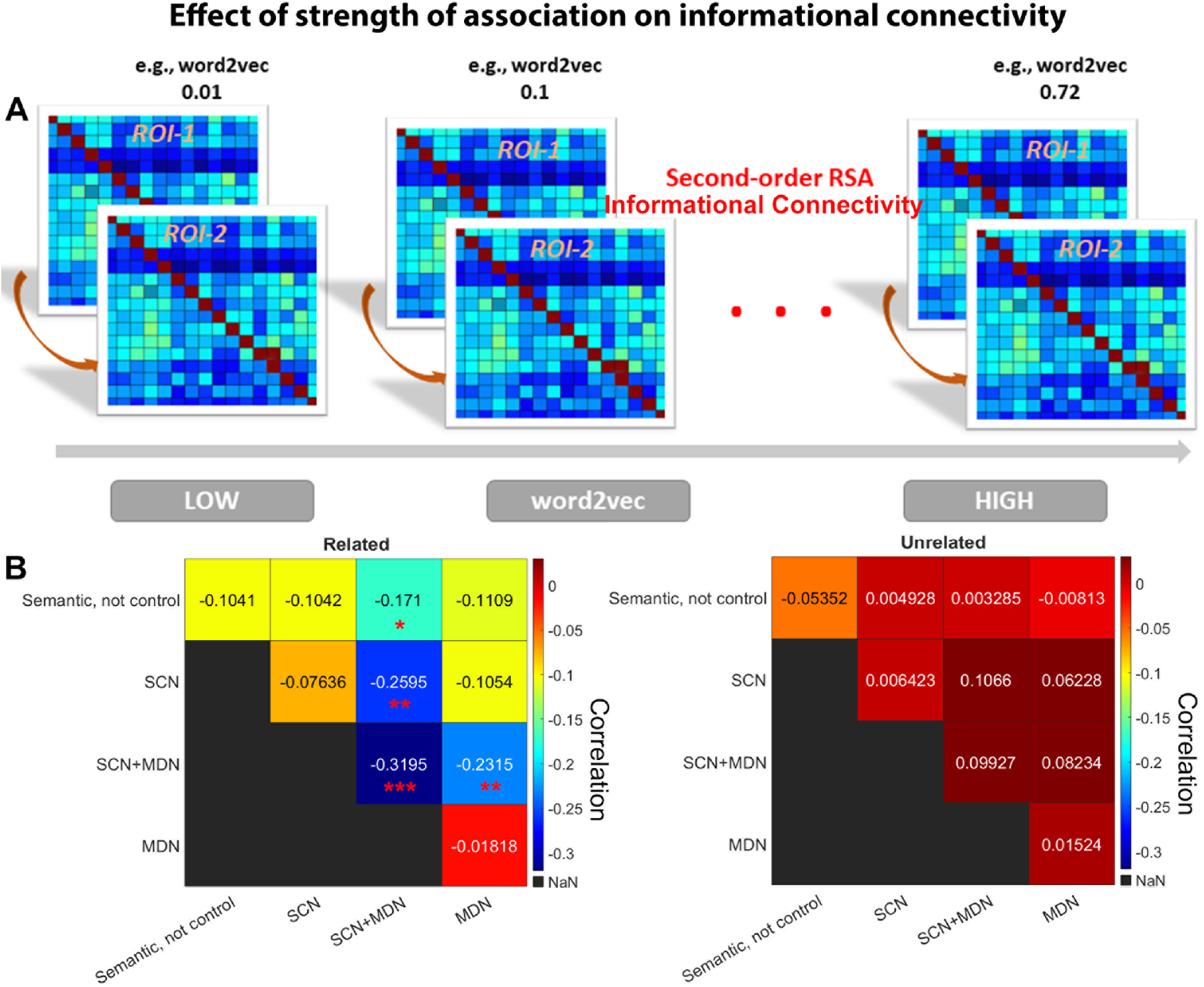
A. Schematic of sliding window analysis of informational connectivity. Trials were sorted according to their association strength from weak to strong associations (based on a word2vec score for each word-pair) and every 16 trials were grouped into one window. We then constructed a neural similarity matrix in each ROI and each window. We measured the informational connectivity within each window by calculating Spearman’s correlation for the neural similarity matrices between ROIs, then averaged across ROIs according to which functional network each site belonged to. Lastly, we calculated a Spearman correlation between association strength and informational connectivity at the network level. B. There was a significant effect of associative strength on informational connectivity between networks for related trials; the multivariate pattern similarity between related trials was increased when strength of association is low for the SCN+MDN regions (left panel); but not for unrelated trials (right panel). * p < 0.05; ** p < 0.01; *** p < 0.001. Bonferroni correction was applied.

To check the robustness of these results, we generated different window sizes containing different numbers of trials along the continuous dimension of association strength, and changed the extent to which adjacent windows overlapped with each other (i.e. the overlap step size). We confirmed the results were robust across a range of window sizes and overlap step sizes (window sizes and overlapping step sizes of 16,4; 12,4; 20,4, respectively). Informational connectivity between SCN+MDN regions and other networks was negatively correlated with association strength in related trials across these analyses (see Supplementary Figure S2B~C).

## Discussion

This study parametrically modulated the association strength between pairs of words to delineate the neural representation of context-free and context-dependent meanings. We related the multivariate neural responses on these trials to two classes of computational linguistic models, representing concepts as either independent or dependent on their linguistic context. Using representational similarity analysis, we found brain activity patterns in the left supramarginal gyrus reflected context-independent conceptual information - but only for the first word that was presented and for trials judged to be semantically unrelated, when there was no linking context to modify the meanings of words. For the second word presented in each pair, there were negative correlations between context-independent semantic models and neural similarity in visual cortex, suggesting that less similar visual features were retrieved for words with similar meanings when participants attempted to retrieve meanings in context. At the same time, context-dependent meanings were represented in regions implicated in semantic control and semantic representation, including left lateral prefrontal cortex and angular gyrus as well as lvATL, on trials judged to be thematically related, when a linking context was retrieved. All large-scale networks implicated in semantic cognition showed this pattern, confirming that the neural response during semantic retrieval tracks the way that words are being interpreted currently (irrespective of associative strength). Despite this network-level similarity, informational connectivity analyses examining multivariate neural similarity across trials found that semantic control regions (defined by the overlap of SCN and MDN) showed more similar patterns across trials to other networks when the words being related were weakly associated. For weak thematic relations, networks were more aligned with control regions, while for strong thematic relations, the responses across networks were more divergent.

Past studies have often compared activation patterns elicited by stimuli from different categories, for instance, faces, objects, places and tools; these studies have significantly advanced our understanding of the neural substrates of ‘individual’ (i.e. static) concepts (Binder et al. 2009; Price 2012). Nevertheless, previous behavioural work on conceptual integration has revealed conceptual representation of word meaning is context sensitive; for instance, when ‘red’ is paired with fire, apple or sky, the magnitude of the representation of ‘red’ is modulated by the following noun (Halff et al. 1976; Coutanche et al. 2019). Previous investigations of dynamic conceptual representation are limited because it is challenging to know how representations of meaning will change between contexts - this information cannot be easily gleaned from participants’ reports. However, ELMo, a recently developed natural language processing algorithm (Peters et al. 2018), allows contextualised conceptual representations to be investigated in the brain. We found context-dependent meaning in all the networks implicated in semantic cognition. Whole-brain analyses also identified distinct clusters in left inferior frontal gyrus within the semantic control network (implicated in controlled semantic retrieval) and left angular gyrus within the default mode network (implicated in more integrative or automatic aspects of semantic retrieval). These effects were only found when semantic links were identified by participants and not when trials were judged to be unrelated. Three recent studies that also employed ELMo and topic modelling techniques to study context-dependent semantic cognition similarly identified left inferior prefrontal and lateral anterior temporal cortex in contextdependent conceptual representation (Lyu et al. 2019; Lopopolo et al. 2020; Toneva et al. 2020). These studies examined the brain’s response to stories and sentences, while our study used a more constrained experimental context which has advantages in terms of experimental control, allowing us to compare neural representations of different decision types and to assess the parametric modulation effect of associative strength on neural representations in a direct and well-controlled fashion.

In a meta-analysis (Binder et al. 2009), left inferior parietal cortex was the region most consistently activated by semantic tasks, but its precise role in semantic cognition is still elusive: it comprises several functionally dissociable areas (Ruschel et al. 2014) and may contribute to both semantic representation and control (Noonan et al. 2013; Humphreys and Lambon Ralph 2015). Our searchlight analysis revealed two clusters in left anterior and posterior lateral parietal cortex, representing context-free and context-dependent meaning respectively. Left SMG showed a positive correlation between neural similarity and context-independent meaning estimated from computational linguistic models. Similarly, a recent RSA study observed that activation patterns in left SMG reflected the semantic similarities of inferred objects (Kivisaari et al. 2019). The anterior cluster within the supramarginal gyrus largely fell within salience and ventral attention networks, which support bottom-up attentional processes (Vossel et al. 2014), and respond to unexpected but salient stimuli (Menon and Uddin 2010; Cai et al. 2019). Decoding using Neurosynth revealed terms linked to attention and cognitive demands. Since left SMG is associated with verbal short-term memory (Buchsbaum and D’Esposito 2009; Baldo et al. 2012), our findings might reflect participants’ need to maintain information about WORD-1 to support the subsequent semantic decision. In contrast, the posterior AG cluster implicated in context-dependent meaning fell within DMN. Decoding using Neurosynth revealed terms linked to semantic memory and language. AG has been linked to the retrieval of thematic knowledge; moreover, this site consistently shows stronger activation to strong than weak associations, implying that it might support more automatic (as well as potentially more controlled) aspects of retrieval (Binder et al. 2009; Humphreys and Lambon Ralph 2015; Jefferies et al. 2020; Humphreys et al. 2021). In line with this, Humphreys and Lambon Ralph (2015) proposed that the inferior parietal lobe (IPL) buffers inputs and learns relations over time, supporting retrieval and integration; however; the time-scales over which it operates may vary from relatively short in anterior IPL (SMG) to longer in posterior IPL (AG). This account might provide an explanation of the functional dissociation we observed in IPL, since SMG might buffer single word inputs (drawing on familiar sequences of phonemes or letters over time), while AG can track semantic contexts given its buffering of more extensive inputs over a long time-period (Lerner et al. 2011; Baldassano et al. 2017).

The control demands of context-dependent meaning retrieval are variable: when words are strongly associated in long-term memory, little control is needed to recover a relevant relationship, since this information is highly accessible. For weak associations, however, recovering a linking context requires controlled retrieval since dominant features and associations not relevant to the linking context must be inhibited. This may help to explain why we observed context-dependent meaning in *both* left IFG (a control site) and AG/lvATL (sites which support more automatic as well as controlled patterns of retrieval). These automatic and controlled aspects of conceptual integration were outside the scope of previous studies using naturalistic stimuli to explore context-dependent meaning (Lyu et al. 2019; Lopopolo et al. 2020; Toneva et al. 2020). Although RSA showed that both control and DMN networks could represent context-dependent meaning irrespective of associative strength, this analysis was blind to potential similarities and differences in the way that context-dependent meaning is represented across trials. Informational connectivity analysis therefore provided complementary evidence. When trials were judged to unrelated, informational connectivity between brain networks was not dependent on the strength of association, remaining relatively stable across windows. A different pattern was found for trials judged to be related: the informational connectivity between networks was more diverse for strong associations as opposed to weak associations, providing evidence that semantic representations coded among regions and networks were different even for the same concepts. Moreover, the multivariate pattern similarity between related trials was higher for weakly-associated items for the SCN+MDN regions, indicating these regions adopt a pattern of connections that supports controlled semantic retrieval. Our results are broadly consistent with the Controlled Semantic Cognition framework that suggests that while a semantic ‘hub’ in ATL might integrate diverse features to form concepts in long-term memory, semantic control regions (both outside and within MDN) might be responsible for supporting the retrieval of non-dominant information required by the context or task instructions (Lambon Ralph et al. 2017). The informational connectivity analysis provided clear evidence that distinct networks played different roles in context-appropriate semantic retrieval.

In previous studies of semantic control, participants have often been asked to focus conceptual retrieval on aspects of knowledge required by the task. For instance, task requirements can gate the recruitment of ‘spoke’ systems (Zhang et al. 2021); participants can retrieve specific unimodal features when they have task instructions providing a clear goal for conceptual processing, and/or suppress activation of non-relevant spoke representations (Coutanche and Thompson-Schill 2014; Martin et al. 2018). In contrast, in the current study, the task instructions did not change between trials: participants were always judging whether two words were thematically related. The meaning of the words themselves defined the nature of the linking context and established which features should be the focus of subsequent retrieval. In this situation, ‘stimulus-driven’ semantic control appears to be supported by the semantic control network, which maintains semantic contexts in a controlled fashion, even when these are non-dominant, to modulate the flow of activation through semantic space. To our best knowledge, this is the first study to compare context-independent and context-dependent meaning representation in the brain during this kind of thematic decision task, which requires meaning-based contexts to drive retrieval.

Left IFG has long been linked to semantic selection and control processes (Thompson-Schill et al. 1997; Jefferies 2013; Noonan et al. 2013; Jackson 2021), and is activated during the retrieval of weak semantic associations (Lambon Ralph et al. 2017; Jefferies et al. 2020). Additional univariate analyses of this dataset focusing on control demands also found higher activation for harder decisions in left IFG and pMTG as well as preSMA (Gao et al. 2021). All of these regions showed successful decoding accuracy of task difficulty, providing strong evidence for their roles in controlled semantic retrieval; however, the contribution of these sites to the representation of conceptual combinations has barely been investigated. One recent study found that left IFG is sensitive to feature uncertainty during the comprehension of combined concepts, while ATL reflects the integration of conceptual features (Solomon and Thompson-Schill 2020). Another recent study investigated how the brain resolves semantic ambiguity in homonym comprehension and found that IFG supports context-appropriate meaning (Hoffman and Tamm 2020). The current study identifies left IFG as one of the sites that supports contextdependent meaning for trials judged to be related - as opposed to context-free meaning for trials judged to be unrelated - implying that left IFG might only represent information suitable for the current context, while inputs that are unable to generate coherent conceptual retrieval might be stored and manipulated in multiple-demand network regions, such as left SMG in the current study.

One limitation of the current study was that our measure of context-sensitive conceptual representation (from ELMo) was derived across trials and participants and was unable to detect individual-specific understanding of each word pair. Moreover, the weaker associations are, the more variance in semantic representation there is likely to be across participants. Future studies could collect subjective reports of context-dependent understanding of word pairs for each participant, and then leverage ELMo to create individual-specific semantic models. More detailed and precise ELMo-based semantic models might result in further neural-semantic alignment results, extending beyond the regions identified here. In addition, we did not find evidence that left ventral ATL represented context-free word meaning in the searchlight analysis, even though this region is thought to provide a heteromodal conceptual ‘hub’ that extracts invariant semantic features across different learning episodes. This site has been shown to decode both the meanings of individual words (Murphy et al. 2017) and contextdependent meaning in previous studies (Lyu et al. 2019; Lopopolo et al. 2020; Toneva et al. 2020). However, ventral parts of ATL are affected by magnetic susceptibility artefacts and our neuroimaging protocol had poorer signal-to-noise in these regions, which may have impacted our ability to resolve neural patterns relating to word meaning. The ROI-based analysis focusing on the lvATL provided evidence for a role of this site in the representation of context-dependent meaning, suggesting future studies using distortion-corrected fMRI techniques may detect stronger effects.

In conclusion, this study leverages natural language models and representational similarity analysis, to compare context-independent and context-dependent meaning representation in the brain during sematic decisions for the first time. Our study demonstrates that different brain regions support context-independent and context-dependent meaning, with a functional dissociation within left IPL between SMG (context-independent representation) and AG (context-dependent representation). In addition, while both regions implicated in relatively automatic (left AG and vATL) and more controlled (left IFG) patterns of semantic retrieval represented context-dependent meaning, the synchronization of neural representation coded in brain networks depended on associative strength, with networks more differentiated from each other as associative strength increased. These findings clarify the roles of distinct brain networks in the computation of coherent meanings across inputs.

## Acknowledgements

This work was sponsored by the European Research Council (Project ID: 771863 - FLEXSEM Project).

## Author contributions

Z.G. and E.J. designed the experiment. K.K.R and X.W. contributed materials. Z.G., A.G., D.V. and X.W. performed the study. L.Z. provided the analysis code. Z.G. analysed the data. Z.G. and E.J. wrote the original manuscript. All authors edited the manuscript.

## References

Aly M, Turk-Browne NB. 2016. Attention promotes episodic encoding by stabilizing hippocampal representations. Proceedings of the National Academy of Sciences. 113:E420–E429.

Andersson JL, Jenkinson M, Smith S. 2007. Non-linear registration aka Spatial normalisation FMRIB Technial Report TR07JA2. FMRIB Analysis Group of the University of Oxford.

Andersson JL, Jenkinson M, Smith S. 2007. Non-linear registration, aka spatial normalisation. FMRIB technial report TR07JA2. 22.

Anzellotti S, Coutanche MN. 2018. Beyond functional connectivity: investigating networks of multivariate representations. Trends in cognitive sciences. 22:258–269.

Baldassano C, Chen J, Zadbood A, Pillow JW, Hasson U, Norman KA. 2017. Discovering event structure in continuous narrative perception and memory. Neuron. 95:709–721. e705.

Baldo JV, Katseff S, Dronkers NF. 2012. Brain regions underlying repetition and auditory-verbal shortterm memory deficits in aphasia: Evidence from voxel-based lesion symptom mapping. Aphasiology. 26:338–354.

Bates D, Mächler M, Bolker B, Walker S. 2014. Fitting linear mixed-effects models using lme4. arXiv preprint arXiv:14065823.

Bellmund JLS, Deuker L, Doeller CF. 2019. Mapping sequence structure in the human lateral entorhinal cortex. eLife. 8:e45333.

Bemis DK, Pylkkänen L. 2013. Basic linguistic composition recruits the left anterior temporal lobe and left angular gyrus during both listening and reading. Cerebral cortex. 23:1859–1873.

Binder JR, Desai RH, Graves WW, Conant LL. 2009. Where is the semantic system? A critical review and meta-analysis of 120 functional neuroimaging studies. Cerebral cortex. 19:2767–2796.

Binney RJ, Embleton KV, Jefferies E, Parker GJM, Lambon Ralph MA. 2010. The Ventral and Inferolateral Aspects of the Anterior Temporal Lobe Are Crucial in Semantic Memory: Evidence from a Novel Direct Comparison of Distortion-Corrected fMRI, rTMS, and Semantic Dementia. Cerebral Cortex. 20:2728–2738.

Binney RJ, Hoffman P, Lambon Ralph MA. 2016. Mapping the Multiple Graded Contributions of the Anterior Temporal Lobe Representational Hub to Abstract and Social Concepts: Evidence from Distortion-corrected fMRI. Cerebral Cortex. 26:4227–4241.

Buchsbaum BR, D’Esposito M. 2009. Repetition Suppression and Reactivation in Auditory–Verbal Short-Term Recognition Memory. Cerebral Cortex. 19:1474–1485.

Cai W, Griffiths K, Korgaonkar MS, Williams LM, Menon V. 2019. Inhibition-related modulation of salience and frontoparietal networks predicts cognitive control ability and inattention symptoms in children with ADHD. Molecular psychiatry.1–10.

Chiou R, Lambon Ralph MA. 2019. Unveiling the dynamic interplay between the hub-and spoke-components of the brain’s semantic system and its impact on human behaviour. NeuroImage. 199:114–126.

Coutanche MN, Solomon S, Thompson-Schill SL. 2019. Conceptual Combination in the Cognitive Neurosciences.

Coutanche MN, Thompson-Schill SL. 2014. Creating Concepts from Converging Features in Human Cortex. Cerebral Cortex. 25:2584–2593.

Davey J, Cornelissen PL, Thompson HE, Sonkusare S, Hallam G, Smallwood J, Jefferies E. 2015. Automatic and controlled semantic retrieval: TMS reveals distinct contributions of posterior middle temporal gyrus and angular gyrus. Journal of Neuroscience. 35:15230–15239.

Davey J, Thompson HE, Hallam G, Karapanagiotidis T, Murphy C, De Caso I, Krieger-Redwood K, Bernhardt BC, Smallwood J, Jefferies E. 2016. Exploring the role of the posterior middle temporal gyrus in semantic cognition: Integration of anterior temporal lobe with executive processes. Neuroimage. 137:165–177.

Deuker L, Bellmund JLS, Navarro Schröder T, Doeller CF. 2016. An event map of memory space in the hippocampus. eLife. 5:e16534.

Fairhall SL, Caramazza A. 2013. Brain Regions That Represent Amodal Conceptual Knowledge. The Journal of Neuroscience. 33:10552–10558.

Fedorenko E, Duncan J, Kanwisher N. 2013. Broad domain generality in focal regions of frontal and parietal cortex. Proceedings of the National Academy of Sciences. 110:16616–16621.

Gao Z, Zheng L, Chiou R, Gouws A, Krieger-Redwood K, Wang X, Varga D, Ralph MAL, Smallwood J, Jefferies E. 2021. Distinct and Common Neural Coding of Semantic and Non-semantic Control Demands. NeuroImage.118230.

Gonzalez Alam TRdJ, Karapanagiotidis T, Smallwood J, Jefferies E. 2019. Degrees of lateralisation in semantic cognition: Evidence from intrinsic connectivity. NeuroImage. 202:116089.

Halff HM, Ortony A, Anderson RC. 1976. A context-sensitive representation of word meanings. Memory & Cognition. 4:378–383.

Hallam GP, Thompson HE, Hymers M, Millman RE, Rodd JM, Ralph MAL, Smallwood J, Jefferies E. 2018. Task-based and resting-state fMRI reveal compensatory network changes following damage to left inferior frontal gyrus. Cortex. 99:150–165.

Hallam GP, Whitney C, Hymers M, Gouws AD, Jefferies E. 2016. Charting the effects of TMS with fMRI: Modulation of cortical recruitment within the distributed network supporting semantic control. Neuropsychologia. 93:40–52.

Hoffman P, Tamm A. 2020. Barking up the right tree: Univariate and multivariate fMRI analyses of homonym comprehension. NeuroImage. 219:117050.

Humphreys GF, Lambon Ralph MA. 2015. Fusion and fission of cognitive functions in the human parietal cortex. Cerebral Cortex. 25:3547–3560.

Humphreys GF, Ralph MAL, Simons JS. 2021. A unifying account of angular gyrus contributions to episodic and semantic cognition. Trends in Neurosciences.

Jackson RL. 2021. The neural correlates of semantic control revisited. NeuroImage. 224:117444.

Jefferies E. 2013. The neural basis of semantic cognition: converging evidence from neuropsychology, neuroimaging and TMS. Cortex. 49:611–625.

Jefferies E, Thompson H, Cornelissen P, Smallwood J. 2020. The neurocognitive basis of knowledge about object identity and events: dissociations reflect opposing effects of semantic coherence and control. Philosophical Transactions of the Royal Society B: Biological Sciences. 375:20190300.

Jung JY, Rice GE, Lambon Ralph MA. 2021. The neural bases of resilient semantic system: evidence of variable neuro-displacement in cognitive systems. Brain Structure and Function. 226:1585–1599.

Kivisaari SL, van Vliet M, Hultén A, Lindh-Knuutila T, Faisal A, Salmelin R. 2019. Reconstructing meaning from bits of information. Nature Communications. 10:927.

Lambon Ralph MA, Jefferies E, Patterson K, Rogers TT. 2017. The neural and computational bases of semantic cognition. Nature Reviews Neuroscience. 18:42.

Lanzoni L, Ravasio D, Thompson H, Vatansever D, Margulies D, Smallwood J, Jefferies E. 2020. The role of default mode network in semantic cue integration. NeuroImage. 219:117019.

Lerner Y, Honey CJ, Silbert LJ, Hasson U. 2011. Topographic Mapping of a Hierarchy of Temporal Receptive Windows Using a Narrated Story. The Journal of Neuroscience. 31:2906–2915.

Leshinskaya A, Contreras JM, Caramazza A, Mitchell JP. 2017. Neural Representations of Belief Concepts: A Representational Similarity Approach to Social Semantics. Cerebral Cortex. 27:344–357.

Lopopolo A, Schoffelen JM, van den Bosch A, Willems RM. 2020. Words in context: tracking context-processing during language comprehension using computational language models and MEG. bioRxiv.2020.2006.2019.161190.

Lyu B, Choi HS, Marslen-Wilson WD, Clarke A, Randall B, Tyler LK. 2019. Neural dynamics of semantic composition. Proceedings of the National Academy of Sciences. 116:21318–21327.

Malone PS, Glezer LS, Kim J, Jiang X, Riesenhuber M. 2016. Multivariate Pattern Analysis Reveals Category-Related Organization of Semantic Representations in Anterior Temporal Cortex. The Journal of Neuroscience. 36:10089–10096.

Martin CB, Douglas D, Newsome RN, Man LL, Barense MD. 2018. Integrative and distinctive coding of visual and conceptual object features in the ventral visual stream. Elife. 7:e31873.

Menon V, Uddin LQ. 2010. Saliency, switching, attention and control: a network model of insula function. Brain structure and function. 214:655–667.

Mumford JA, Poldrack RA. 2007. Modeling group fMRI data. Social cognitive and affective neuroscience. 2:251–257.

Murphy C, Rueschemeyer S-A, Watson D, Karapanagiotidis T, Smallwood J, Jefferies E. 2017. Fractionating the anterior temporal lobe: MVPA reveals differential responses to input and conceptual modality. NeuroImage. 147:19–31.

Noonan KA, Jefferies E, Visser M, Lambon Ralph MA. 2013. Going beyond inferior prefrontal involvement in semantic control: evidence for the additional contribution of dorsal angular gyrus and posterior middle temporal cortex. Journal of cognitive neuroscience. 25:1824–1850.

Peters ME, Neumann M, Iyyer M, Gardner M, Clark C, Lee K, Zettlemoyer L. 2018. Deep contextualized word representations. arXiv preprint arXiv:180205365.

Price CJ. 2012. A review and synthesis of the first 20 years of PET and fMRI studies of heard speech, spoken language and reading. Neuroimage. 62:816–847.

Ruschel M, Knösche TR, Friederici AD, Turner R, Geyer S, Anwander A. 2014. Connectivity architecture and subdivision of the human inferior parietal cortex revealed by diffusion MRI. Cerebral cortex. 24:2436–2448.

Solomon SH, Thompson-Schill SL. 2020. Feature Uncertainty Predicts Behavioral and Neural Responses to Combined Concepts. The Journal of Neuroscience. 40:4900–4912.

Stolier RM, Freeman JB. 2016. Neural pattern similarity reveals the inherent intersection of social categories. Nature Neuroscience. 19:795–797.

Teige C, Cornelissen PL, Mollo G, Gonzalez Alam TRDJ, Mccarty K, Smallwood J, Jefferies E. 2019. Dissociations in semantic cognition: Oscillatory evidence for opposing effects of semantic control and type of semantic relation in anterior and posterior temporal cortex. Cortex. 120:308–325.

Thompson-Schill SL, D’Esposito M, Aguirre GK, Farah MJ. 1997. Role of left inferior prefrontal cortex in retrieval of semantic knowledge: a reevaluation. Proceedings of the National Academy of Sciences. 94:14792–14797.

Toneva M, Mitchell TM, Wehbe L. 2020. Combining computational controls with natural text reveals new aspects of meaning composition. bioRxiv.2020.2009.2028.316935.

Viganò S, Piazza M. 2020. Distance and Direction Codes Underlie Navigation of a Novel Semantic Space in the Human Brain. The Journal of Neuroscience. 40:2727–2736.

Vossel S, Geng JJ, Fink GR. 2014. Dorsal and ventral attention systems: distinct neural circuits but collaborative roles. The Neuroscientist. 20:150–159.

Wagner AD, Paré-Blagoev EJ, Clark J, Poldrack RA. 2001. Recovering meaning: left prefrontal cortex guides controlled semantic retrieval. Neuron. 31:329–338.

Wang X, Margulies DS, Smallwood J, Jefferies E. 2020. A gradient from long-term memory to novel cognition: Transitions through default mode and executive cortex. NeuroImage. 220:117074.

Wang X, Wu W, Ling Z, Xu Y, Fang Y, Wang X, Binder JR, Men W, Gao J-H, Bi Y. 2017. Organizational Principles of Abstract Words in the Human Brain. Cerebral Cortex. 28:4305–4318.

Ward EJ, Chun MM, Kuhl BA. 2013. Repetition suppression and multi-voxel pattern similarity differentially track implicit and explicit visual memory. Journal of Neuroscience. 33:14749–14757.

Weiskopf N, Hutton C, Josephs O, Deichmann R. 2006. Optimal EPI parameters for reduction of susceptibility-induced BOLD sensitivity losses: a whole-brain analysis at 3 T and 1.5 T. Neuroimage. 33:493–504.

Whitney C, Kirk M, O’Sullivan J, Lambon Ralph MA, Jefferies E. 2011. The neural organization of semantic control: TMS evidence for a distributed network in left inferior frontal and posterior middle temporal gyrus. Cerebral cortex. 21:1066–1075.

Xiao X, Dong Q, Gao J, Men W, Poldrack RA, Xue G. 2017. Transformed neural pattern reinstatement during episodic memory retrieval. Journal of Neuroscience. 37:2986–2998.

Xu Y, Wang X, Wang X, Men W, Gao J-H, Bi Y. 2018. Doctor, Teacher, and Stethoscope: Neural Representation of Different Types of Semantic Relations. The Journal of Neuroscience. 38:3303–3317.

Zhang M, Varga D, Wang X, Krieger-Redwood K, Gouws A, Smallwood J, Jefferies E. 2021. Knowing what you need to know in advance: The neural processes underpinning flexible semantic retrieval of thematic and taxonomic relations. NeuroImage. 224:117405.

