## Supplementary Materials for "Context Free and Context-Dependent Conceptual Representation in the Brain"

**Supplementary Material**

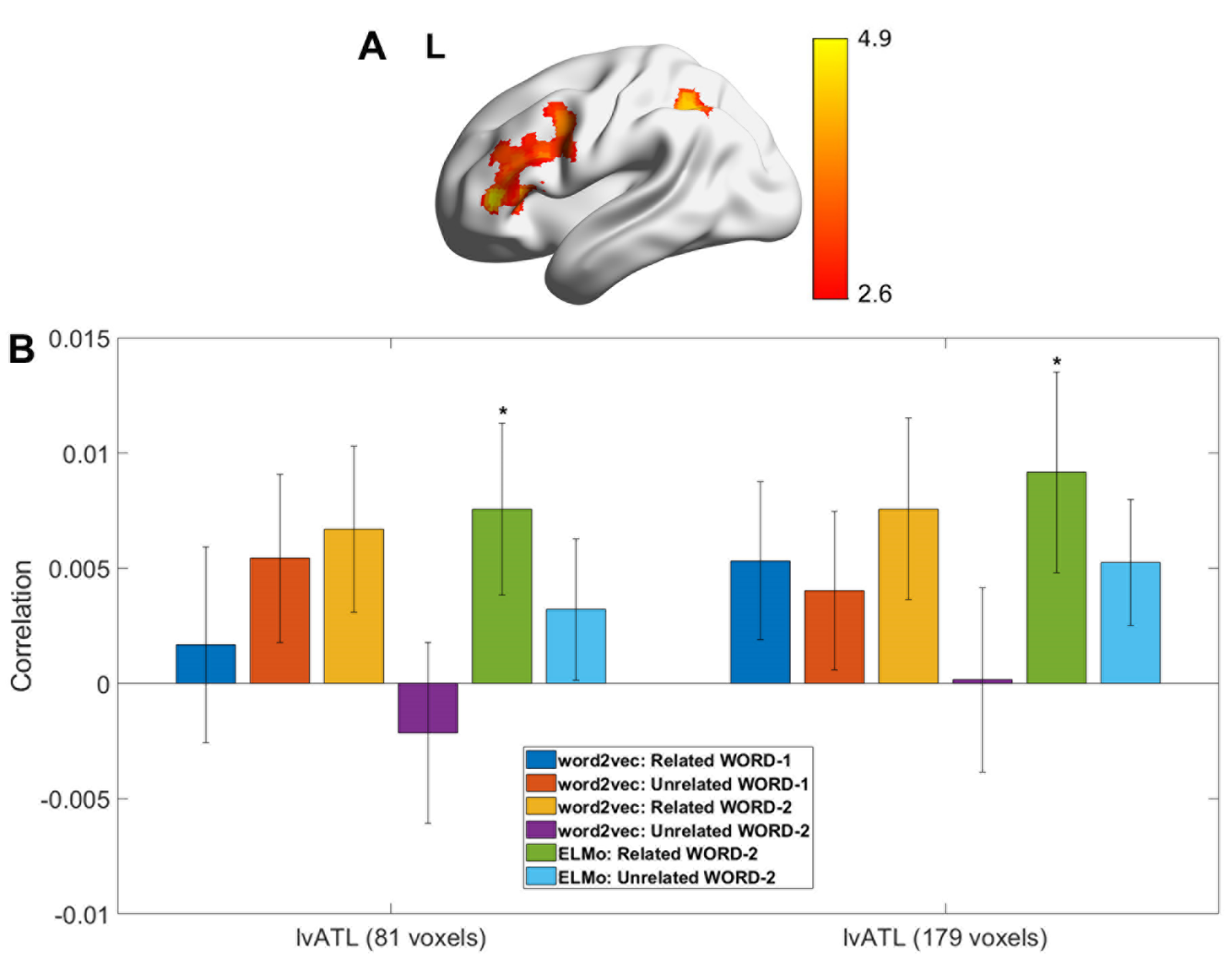

Figure S1. A. Neural representation of the context-dependent meaning of WORD-2 in trials judged to be related. The context-dependent meaning RSM was derived from sentence-based ELMo embedding vectors, in contrast to the findings in the main body which used embedding vectors based on pairs of words (Z > 2.6, corrected). B. Neural representation of context-free and context-dependent meaning in left ventral anterior temporal lobe (lvATL) for two sphere sizes. Positive correlations were only found for context-dependent meanings of WORD-2 on trials judged to be related. * p < 0.05

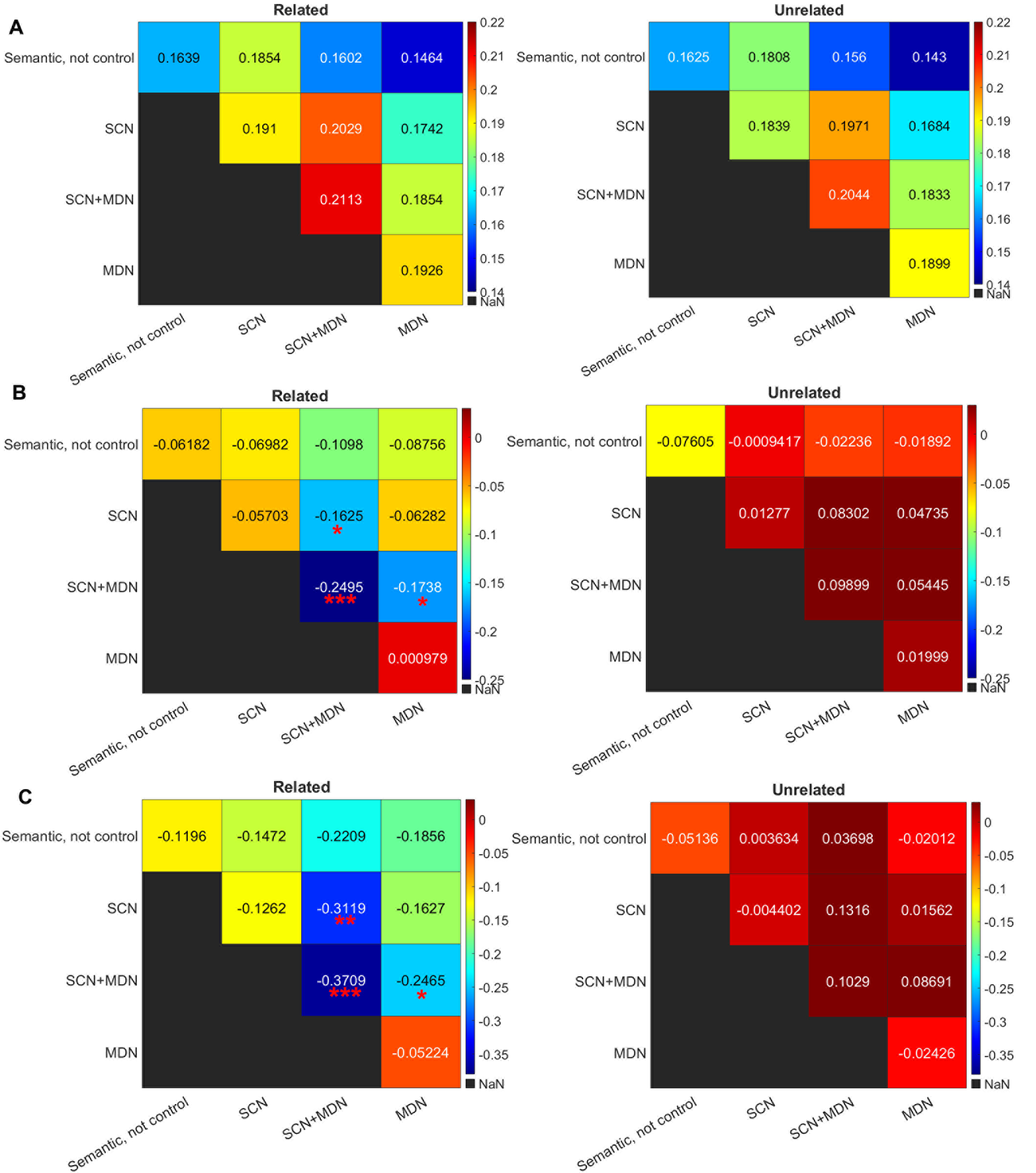

Figure S2. A. Overall informational connectivity when all trials were included (examining trials judged to be related and unrelated separately). All within network and between network informational connectivity metrics were significantly higher than chance level (i.e., greater than zero; p < 0.001 with Bonferroni correction). Direct comparisons of informational connectivity for related and unrelated trials found no significant differences (all Ps > 0.23 before multiple comparison correction. B and C. Sliding window informational connectivity examining the effects of strength of association, using two different window sizes. B. Window length: 12 trials, step size: 4 trials; C. Window length: 20 trials, step size, 4 trials. There was a significant effect of associative strength on informational connectivity between networks for trials judged to be related; the multivariate pattern similarity between related trials was increased when strength of association was low within SCN+MDN regions (left panel); but this effect was not found for trials judged to be unrelated (right panel). SCN = semantic control network. SCN+MDN = multiple-demand regions implicated in semantic control. MDN = multiple-demand regions not implicated in semantic cognition. * p < 0.05; ** p < 0.01; *** p < 0.001, with Bonferroni correction applied.

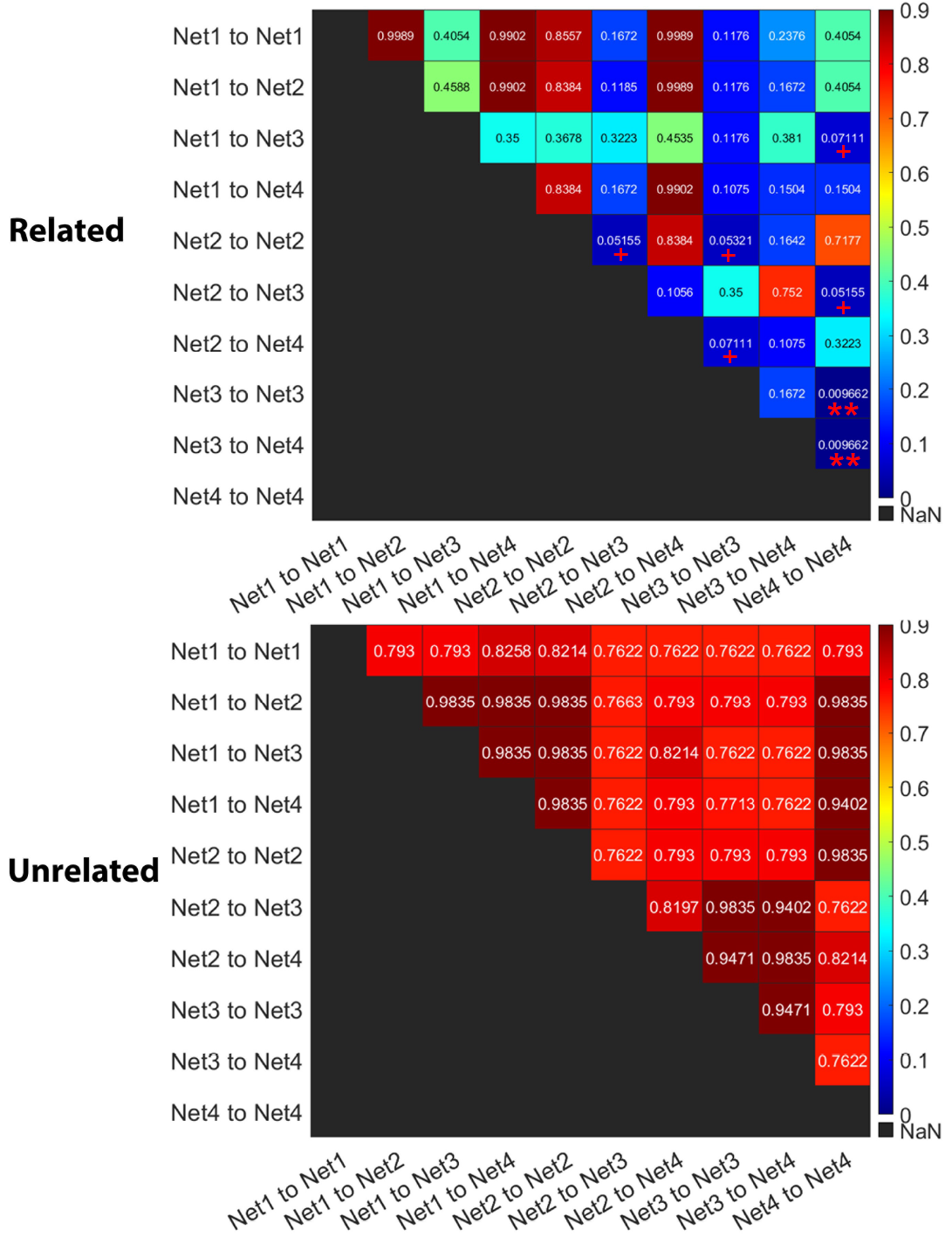

Figure S3. Direct comparisons of the modulation effects of associative strength on informational connectivity for trials judged to be related (left panel) and unrelated (right panel), when pairs of networks are compared. p values in each grid are after the application of multiple comparisons correction using FDR (Benjamini-Hochberg). SCN+MDN is implicated in every grid that shows significant or marginal significant effect. Net1: Semantic, not control; Net2, SCN; Net3, SCN+MDN; Net4, MDN.

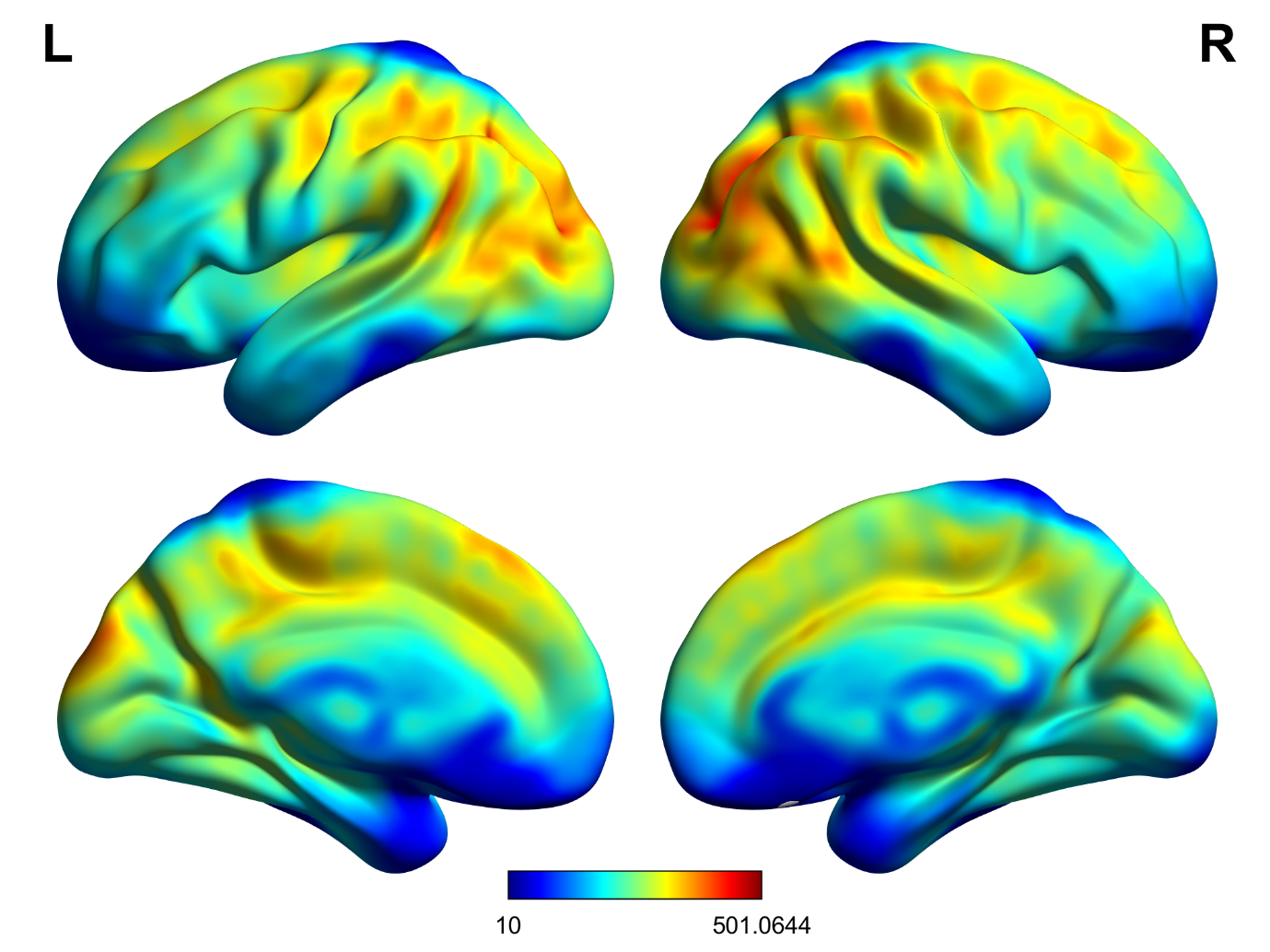

Figure S4. tSNR map. We calculated the temporal signal-to-noise ratio (tSNR) for each participant by dividing the mean of the smoothed time series in each voxel by its standard deviation in each run; we then averaged the tSNR across all runs for the semantic task. These tSNR values were comparable with previous studies (Hoffman et al. 2015; Striem-Amit et al. 2018), and at an acceptable level (Murphy et al. 2007), though lowest at the anterior temporal pole (mean value: 107.8). The full tSNR map in MNI space is available to view online: https://neurovault.org/images/441927/.”

Hoffman P, Binney RJ, Ralph MAL. 2015. Differing contributions of inferior prefrontal and anterior temporal cortex to concrete and abstract conceptual knowledge. Cortex. 63:250-266.

Murphy K, Bodurka J, Bandettini PA. 2007. How long to scan? The relationship between fMRI temporal signal to noise ratio and necessary scan duration. Neuroimage. 34:565-574.

Striem-Amit E, Wang X, Bi Y, Caramazza A. 2018. Neural representation of visual concepts in people born blind. Nature Communications. 9:5250.
